# Right-Lateralized Maladaptive Topological Reorganization in Hyperthyroidism

**DOI:** 10.1101/2025.10.16.682796

**Authors:** Priyanka Chakraborty, Neeraj Upadhyay, Pawan Kumar, Poonam Rana, Suman Saha, Maria D’souza, Tarun Sekhri, Subash Khushu, Mukesh Kumar

## Abstract

Hyperthyroidism (HT) has been associated with cognitive impairments, but the structural connectomic and molecular mechanisms underlying these deficits remain poorly understood. In this study, we investigated large-scale brain network reorganization using structural connectivity derived from diffusion MRI in patients with 30 hyperthyroidism and 28 matched control participants. Subject level connectomes were analyzed to characterize network topology, with a specific focus on hub architecture, nodal alterations, and hemispheric reorganization. To probe molecular underpinnings, we integrated normative neurotransmitter receptor and transporter density maps with network metrics, assessing their correspondence to observed connectomic disruptions.

Individuals with hyperthyroidism exhibited increased structural connectivity, particularly in mid- and long-range connections, alongside disrupted network topology. These changes were characterized by reduced modularity, increased characteristic path length and hub reorganization, most prominently in subcortical and right-hemisphere regions. Importantly, network alterations were functionally relevant, as metrics showed significant associations with clinical and cognitive variables, including FT4 levels, BMI, and perceptuomotor performance in the hyperthyroid group. Spatial neuromolecular correspondence analyses further revealed that DAT, 5-HT2A, and NET were closely aligned with topological changes. Moreover, neurotransmitter-weighted within-module degree z-scores moderately predicted TSH levels in HT.

Our findings demonstrate that hyperthyroidism is associated with an increase in redundant, spatially extended white matter connections and a marked right-lateralized asymmetry in network organization. Furthermore, serotonergic (5-HT1a) and dopaminergic (DAT) systems consistently contributed to the modulation of intra-modular hubness, suggesting their role in maladaptive structural reorganization that may underlie perceptuomotor dysfunction in hyperthyroidism.

## 1. Introduction

Triiodothyronine (T3) and free thyroxine (fT4) are thyroid hormones (THs) essential for neuronal differentiation, neurogenesis, and myelination throughout life (Bernal et al., 2003; Calzà et al., 2015). Elevated T3 and/or fT4 levels with suppressed thyroid-stimulating hormone (TSH) define hyperthyroidism (HT) (Lee & Pearce, 2023), a condition characterized by increased metabolism, weight loss, tremors, anxiety, depression, and cardiovascular irregularities (Chiera et al., 2023; Dumitriu & Ursu, 1985; Mattesi et al., 2022). Recent evidence links TSH to metabolic regulation via brain–body communication (Wang et al., 2024), and cognitive deficits involving memory, decision-making, and emotion regulation are well-documented in HT (Eslami-Amirabadi & Sajjadi, 2021; Holmberg et al., 2019). Meta-analytic data suggest that HT increases dementia risk (Ma et al., 2023). Structurally, higher fT4 levels correlate with larger intracranial, brain, and white matter volumes in younger adults (Chaker et al., 2017), indicating that TH dysfunction may alter brain architecture, leading to abnormal metabolism, mood disturbances, and cognitive decline. Thus, examining structural reorganization in HT is crucial for understanding its pathophysiology.

Diffusion-weighted imaging (DWI) enables characterization of white matter integrity and structural connectivity (Mori & Zhang, 2006). Studies in thyroid-associated ophthalmopathy (TAO) have reported reduced fractional anisotropy and axial diffusivity, along with increased mean and radial diffusivity, particularly in the right occipital and cuneus regions, indicating microstructural alterations (Wu et al., 2020; Yeo et al., 2025). Graph-theoretical analyses have revealed preserved global but altered nodal network properties in TAO (Wu et al., 2021), including decreased nodal degree and efficiency in prefrontal and cingulate areas and increased degree in the left cuneus. These results suggest compensatory reorganization, but how structural networks reorganize under overt HT to maintain or disrupt function remains unknown.

Beyond structural changes, HT is associated with alterations in dopaminergic, noradrenergic, serotonergic, and GABAergic neurotransmission (Napoli et al., 2001; Rastogi & Singhal, 1976; Wiens & Trudeau, 2006a). Elevated serotonin (5-HT) levels in cortical and brainstem regions indicate that thyroid hormones modulate mood and behavior through region-specific monoaminergic pathways (Bauer et al., 2002; Ito et al., 1977). Enhanced dopamine accumulation in the mesodiencephalon further supports altered decision-making in HT (Bauer et al., 2002). The neuromodulator 5-HT also shapes cytoarchitecture and connectivity during development (Daubert & Condron, 2010), and regional variations in neurotransmitter densities have been linked to structural network organization (Hansen et al., 2022). Hence, imbalances in neurotransmitter systems may underlie the cognitive and behavioral disturbances observed in HT.

Despite evidence of structural and neurochemical abnormalities, the connectomic mechanisms linking thyroid hormone dysregulation to brain network organization and cognitive deficits remain poorly understood. This study therefore aims to:(1) Characterize structural connectome reorganization in hyperthyroidism, emphasizing global topology, hub configuration, and hemispheric asymmetry; (2) Examine associations between altered network measures, thyroid hormone levels, and cognitive performance; and (3) Link macroscale connectomic changes to the normative spatial distribution of neurotransmitter receptor and transporter densities. By integrating structural connectivity, endocrine, cognitive, and molecular data, this multiscale approach provides new insights into the mechanisms of brain network vulnerability and cognitive dysfunction in hyperthyroidism.

## 2. Materials and methods

### 2.1 Subjects

Thirty drug-naive hyperthyroid patients (34.63±8.99 years; 10M/20F) and 28 healthy controls (32.82±9.86 years; 9M/19F) were recruited. Patients were newly diagnosed with clinical symptoms including palpitation, tremor, weight loss, and heat intolerance. Exclusion criteria included trauma, psychiatric illness, substance abuse, pregnancy, or MRI contraindications. Hyperthyroid patients were recruited from the Thyroid Research Centre, INMAS, India, and controls from the local community. The study was approved by the institutional ethics committee (IIHEC/CT/2017/15), and written informed consent was obtained.

### 2.2 Clinical assessment/Thyroid Hormone tests

Blood samples were collected in the morning under fasting conditions. Serum TSH, free T4 (fT4), and free T3 (fT3) were measured using an electrochemiluminescence immunoassay (Elecsys 2010; Roche Diagnostics, Mannheim, Germany). Controls had hormone levels within normal ranges, whereas hyperthyroid patients showed suppressed TSH (0.01±0.019 µIU/ml) and elevated fT4 (42.91±18.58 pmol/l) and fT3 (15.46±5.56 pmol/l) (Table 1).

**Table 1:**
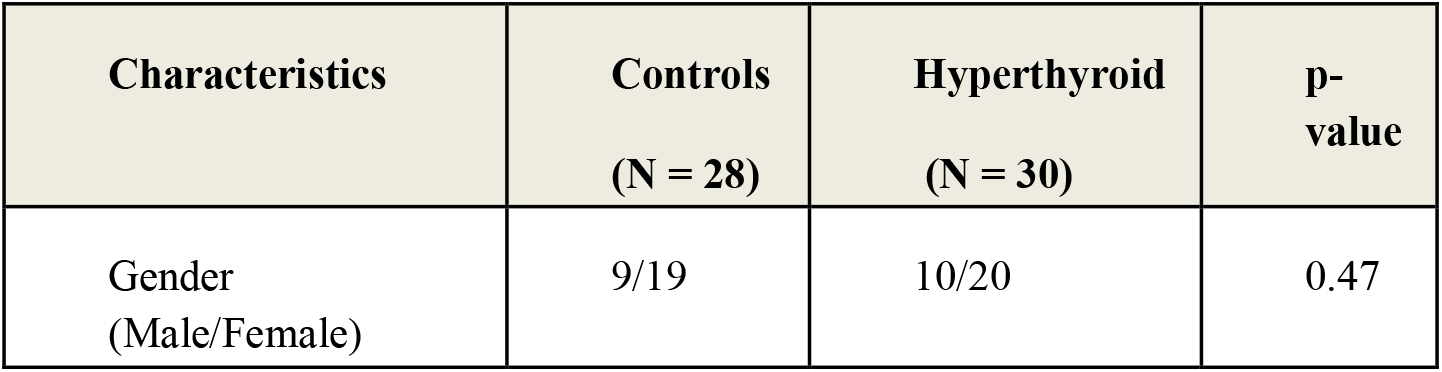
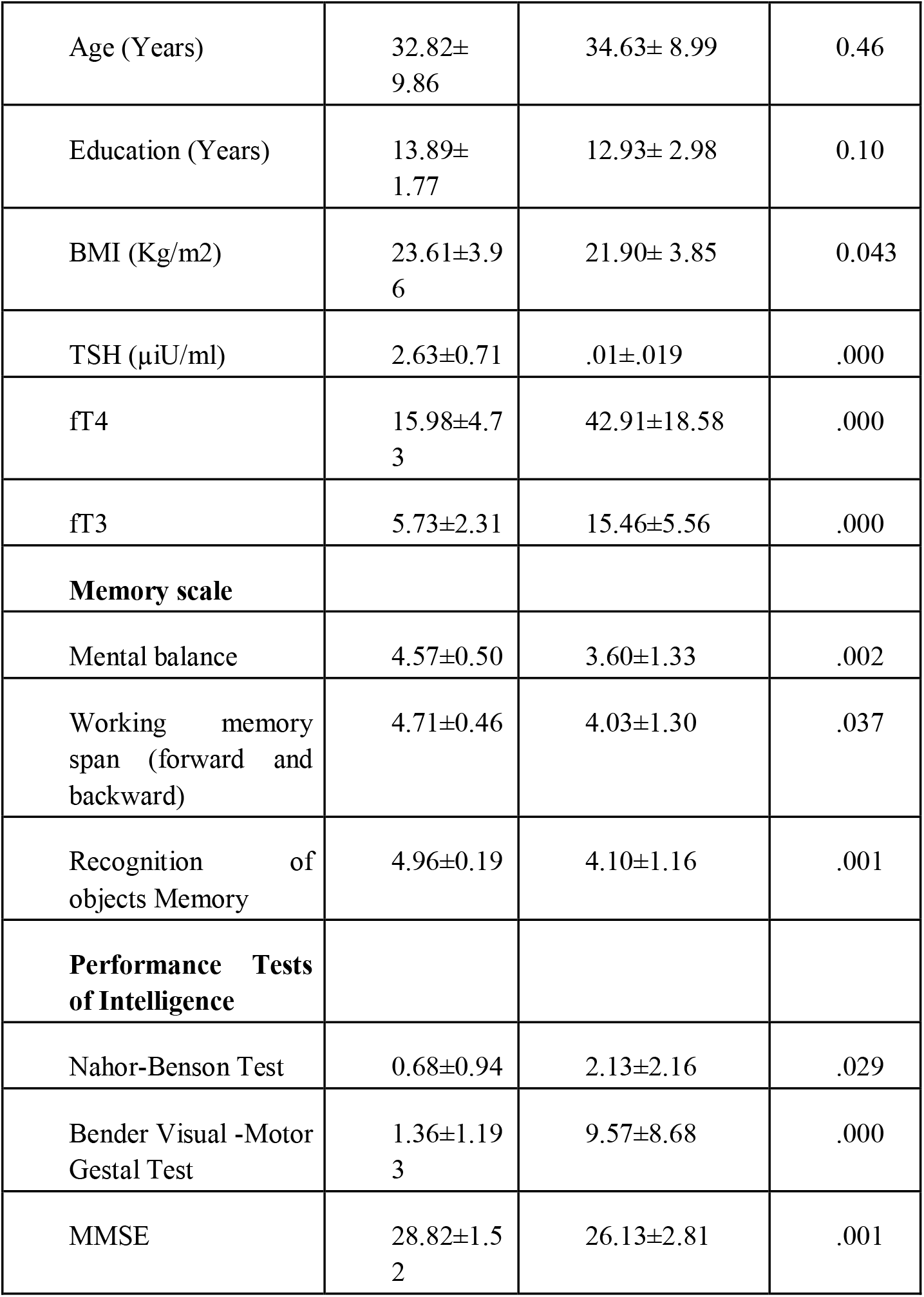
Demographic, clinical and neuropsychological characteristics of hyperthyroid, as compared to healthy controls.

| Characteristics | Controls<br>(N = 28) | Hyperthyroid<br>(N = 30) | p-value |
| --- | --- | --- | --- |
| Gender<br>(Male/Female) | 9/19 | 10/20 | 0.47 |
| Age (Years) | 32.82±<br>9.86 | 34.63± 8.99 | 0.46 |
| Education (Years) | 13.89±<br>1.77 | 12.93± 2.98 | 0.10 |
| BMI (Kg/m2) | 23.61±3.9<br>6 | 21.90± 3.85 | 0.043 |
| TSH (µIU/ml) | 2.63±0.71 | .01±.019 | .000 |
| fT4 | 15.98±4.7<br>3 | 42.91±18.58 | .000 |
| fT3 | 5.73±2.31 | 15.46±5.56 | .000 |
| **Memory scale** |  |  |  |
| Mental balance | 4.57±0.50 | 3.60±1.33 | .002 |
| Working memory span (forward and backward) | 4.71±0.46 | 4.03±1.30 | .037 |
| Recognition of objects Memory | 4.96±0.19 | 4.10±1.16 | .001 |
| **Performance Tests of Intelligence** |  |  |  |
| Nahor-Benson Test | 0.68±0.94 | 2.13±2.16 | .029 |
| Bender Visual -Motor Gestal Test | 1.36±1.19<br>3 | 9.57±8.68 | .000 |
| MMSE | 28.82±1.5<br>2 | 26.13±2.81 | .001 |

**Table 2:** Comparison of network metrics between HC and HT. Values are reported as mean ± standard deviation. Cohen’s d indicates the effect size of the group difference. Positive Cohen’s d indicates higher values in HT compared to HC, while negative values indicate lower values in HT.

| Metric | HC (Mean ± SD) | HT (Mean ± SD) | p-value | Cohen's d |
| --- | --- | --- | --- | --- |
| Total SC count | $1.69 \times 10^9 \pm 2.91 \times 10^8$ | $1.81 \times 10^9 \pm 1.94 \times 10^8$ | 0.065 | 0.49 |
| SR count | $7.73 \times 10^8 \pm 1.27 \times 10^8$ | $8.08 \times 10^8 \pm 8.66 \times 10^7$ | 0.221 | 0.33 |
| MR count | $6.90 \times 10^7 \pm 3.35 \times 10^7$ | $9.30 \times 10^7 \pm 2.42 \times 10^7$ | 0.002 | 0.83 |
| LR count | $2.30 \times 10^6 \pm 1.85 \times 10^6$ | $3.77 \times 10^6 \pm 2.63 \times 10^6$ | 0.018 | 0.64 |
| Modularity | $0.58 \pm 0.03$ | $0.56 \pm 0.02$ | 0.009 | -0.71 |
| Char. Path length | $0.28 \pm 0.08$ | $0.32 \pm 0.04$ | 0.032 | 0.58 |
| Global efficiency | $0.05 \pm 0.01$ | $0.05 \pm 0.01$ | 0.156 | 0.38 |
| Transitivity | $0.001 \pm 0.002$ | $0.001 \pm 0.003$ | 0.811 | 0.06 |

**Table 3:** Association between cognitive and clinical scores with global network measures for HT group.

| Cognitive/ clinical | Network measures | Coefficient | R2 | p |
| --- | --- | --- | --- | --- |
| BMI | Modularity | 1.12 | 0.21 | 0.04 |
| FT4 | Char. path length | 0.392 | 0.17 | 0.01 |
| FT4 | LR connection strength | 0.376 | 0.14 | 0.03 |

**Table 4:** Association between cognitive scores with local network measures for HT group.

| Cognitive/ clinical | Network measures | ROI | Coefficient | R2 | p |
| --- | --- | --- | --- | --- | --- |
| Nahor-Benson Test | MR connection strength | Right Cerebelum IV-V | 3.706 | 0.143 | 0.02 |
| Nahor-Benson Test | Degree | Right Cerebelum IV-V | 1.82 | 0.119 | 0.03 |

### 2.3 Neuropsychological Data

All participants completed the Mini-Mental State Examination (MMSE) (Folstein et al., 1975) and the Postgraduate Institute Battery of Brain Dysfunction (PGIBBD) (Pershad, 1990), assessing memory, executive, motor, and visuospatial functions. Higher scores indicated better performance on memory and executive tasks, whereas higher dysfunction scores reflected poorer motor or visuospatial performance (Pershad & Verma, 1990).

### 2.4 Structural data acquisition

MRI was performed on a 3T Siemens Magnetom Skyra with a 20-channel head coil. Diffusion-weighted images were acquired using a single-shot echo-planar dual SE sequence with multiple b-values (0–2000 s/mm^2^) and 30 diffusion directions. Routine T1- and T2-weighted images were obtained to rule out structural abnormalities.

### 2.5 DTI preprocessing

The diffusion MRI data were processed locally using our pre-processing pipeline, which involved using MRtrix3 ((Tournier et al.2019), https://github.com/MRtrix3/mrtrix3), FSL (http://www.fmrib.ox.ac.uk/fsl), and ANTs (http://stnava.github.io/ANTs/). The main preprocessing steps included described in supplementary material.

#### Structural connectivity and fiber tract length

Whole-brain connectomes were generated using the AAL atlas (Tzourio-Mazoyer et al., 2002), calculating fiber density and tract length (≤250 mm) between each ROI. Connections were classified as short, medium, or long (70 and 140 mm thresholds) (Chakraborty et al., 2025). A group consensus network preserving edge-density and length distributions was created, with weights averaged across participants and scaled 0-1 (Betzel et al., 2019).

#### Graph theoretical network metrics

Node- and network-level metrics, including degree, clustering coefficient (CC), within-module degree (WD), betweenness centrality (BW), participation coefficient (PC), modularity, transitivity, characteristic path length, and global efficiency, were computed using the Brain Connectivity Toolbox (Rubinov & Sporns, 2010).

#### Structural core hub detection algorithm

We classified the 116 brain regions into three main categories: non-hubs, provincial hubs, and connector hubs.

1. We first applied a consensus-based approach on both groups and in average matrix as well as subject wise matrix.
2. We then identified nodes whose degree exceeded the group-specific mean by more than one standard deviation.
3. Among the remaining nodes, we selected the top 50% with the highest betweenness centrality(BW).
4. Nodes within module degree z-score(WD), WD>1 were classified as hubs, while those with WD<1 were considered non-hubs (Fig. S3a)
5. Finally, among the identified hubs, nodes with a participation coefficient (PC), PC<0.3 were labeled provincial hubs, and those with PC>0.3 were labeled connector hubs (Guimerà & Nunes Amaral, 2005).

### 2.6 Neurotransmitter maps from positron emission tomography

Regional densities for 19 receptors/transporters across nine neurotransmitter systems were extracted using PET studies (Hansen, Shafiei, Markello, et al., 2022), including dopamine (DAT), norepinephrine (NET), serotonin (5-HT), and GABA (Napoli et al., 2001; Rastogi & Singhal, 1976; Wiens & Trudeau, 2006b). We extracted regional neurotransmitter densities for these 9 receptors and transporters using the AAL atlas, then z-scored each map before compiling them into a region × receptor matrix of relative densities.

### 2.7 Neurotransmitter-weighted network (NN) features

For each subject, different nodal graph-theoretical metrics-degree, CC, BW, WD, and PC-were correlated with the nine neurotransmitter density maps, resulting in a 30×9 neurotransmitter-weighted network (NN) feature matrix for each metric in the HT group (and 28×9 for the HC group).

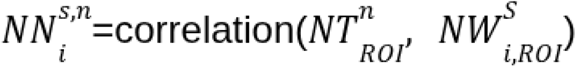

where NT^n^_ROI_ is the ROI-wise density of neurotransmitter n, and NW_i,ROIS_ is the ROI-wise value of the i-th network metric for subjects. Here, I represent network measures as degree, CC, BW, WD, or PC. Here, we chose to exclude the vermis regions (ROI IDs 109-116), as spin permutation was applied using hemispheric rotations on each hemisphere separately (Hansen et al., 2022).

### 2.8 Statistical analysis

Demographic and clinical comparisons used t-tests or Chi-square; ANCOVA for neuropsychological scores (covariates: age, gender). Brain network differences were assessed using t-tests with FDR correction (Benjamini & Hochberg, 1995) and permutation testing (n=10,000). Spin permutation tests accounted for spatial autocorrelation in receptor maps (Alexander-Bloch et al., 2018; Markello & Mišić, 2021). Multiple linear regression and dominance analysis (Azen & Budescu) were applied to evaluate predictors of network measures, controlling for age, gender, and education, with generalizability tested via 10,000 permutations.

## 3. Results

### 3.1 Clinical and Demographic results

Demographic, clinical, and neuropsychological scores are summarized in Table 1. Groups did not differ in age (p=0.46), gender (p=0.47), or education (p=0.10), but BMI was lower in hyperthyroid patients (p=0.043). Hyperthyroid patients showed lower MMSE (p=0.001), mental balance (p=0.002), working memory (p=0.037), immediate semantic recall (p=0.05), object recognition (p=0.001), Bender Gestalt (p<0.001), and Nahar-Benson (p=0.029) scores compared to controls. They also exhibited suppressed TSH (p<0.001) and elevated fT3 (p<0.001) and fT4 (p<0.001) levels.

Mean and standard deviation (SD) for demographic, blood hormone levels, neuropsychological tests score.

### 3.2 Structural topology alterations in hyperthyroidism

We first assessed whole-brain structural connectivity (SC) patterns in individual subjects from the HC and HT groups. Representative SC matrices illustrated widespread structural connections in both groups (Fig. 1a-b). We then divide the structural connections based on their physical length. Distribution of tract length for control group shown in Fig. 1c. Total connection strength remains the same for two groups (Fig. 1d left). We checked that short-range (SR) connections count/strength showed a non-significant trend toward being higher in the HT group compared to the HC group. In contrast, both medium-range (MR) and long-range (LR) connections strength were significantly elevated in the HT group (MR: p = 0.005; LR: p = 0.012, Fig. 1d). Mean and standard division of each metric were presented in table2.

**Figure 1.**
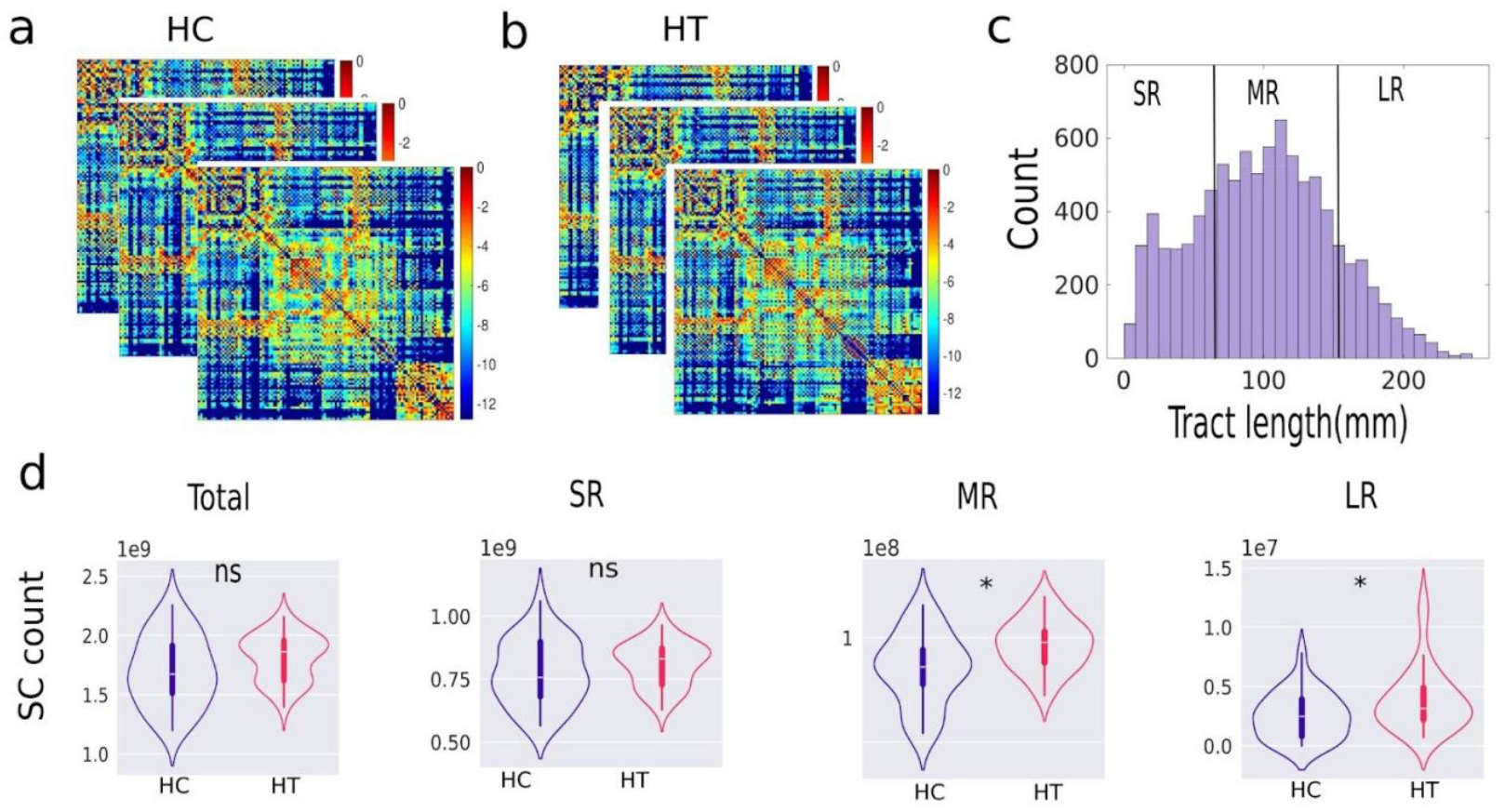
Group differences in structural connectivity patterns between control (HC) and hyperthyroid (HT) groups. (a-b) Representative structural connectivity matrices from individual subjects in the HC (a) and HT (b) groups. (c)Tract length distribution for the HC group, with vertical lines indicating the lower (70 mm) and upper (140 mm) thresholds used to define connection categories. Connections are classified as short-range (SR), medium-range (MR), or long-range (LR) based on these thresholds. (d) Count/strength of total connections, SR, MR, and LR connections for the two groups are shown, respectively. Asterisks indicate significant differences (p < 0.05); ‘ns’ denotes non-significant comparisons.

**Figure 2.**
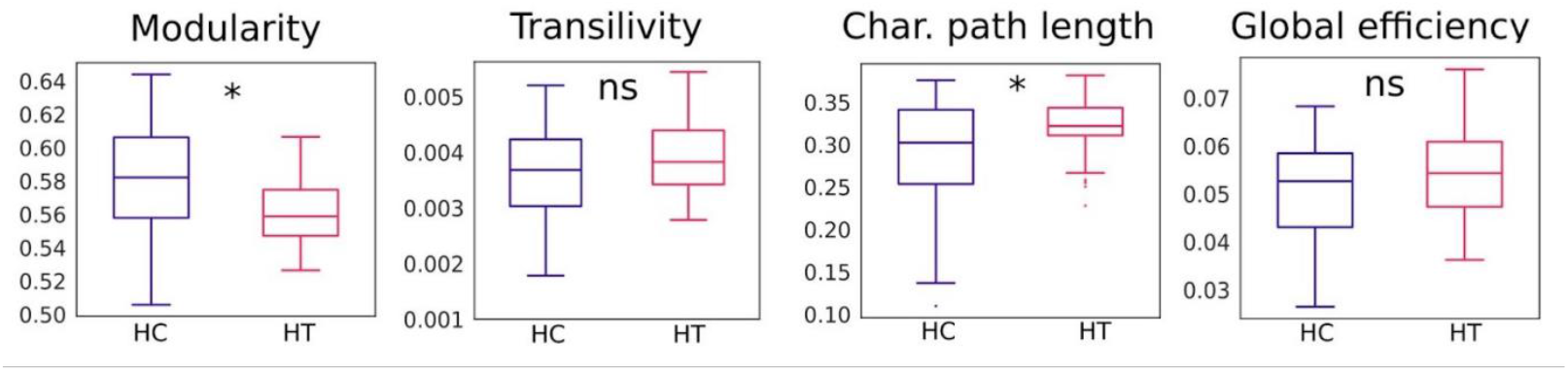
Boxplots comparing global network measures between groups over subjects. Modularity and characteristic path length is significantly reduced and increased in the HT group, (p = 0.01 and p=0.03) respectively. No significant (ns) group differences were observed for transitivity and global efficiency.

#### Global graph measures in HT

We further examined global structural network organization in the HC and HT groups. Subject-level comparisons of global graph metrics revealed a significant reduction in modularity while an increase in char path length in the HT group compared to HC (p < 0.05), indicating decreased structural segregation and information efficiency. No significant group differences were observed in transitivity and global efficiency.

#### Regional changes in HT

Subject-level comparisons revealed significant topological alterations across multiple regions in both hemispheres. In the left hemisphere, the hyperthyroid (HT) group showed higher nodal degree (FDR-corrected, p < 0.05) in widespread cortical, subcortical, and cerebellar areas, including the paracentral lobule, supplementary motor area, insula, cingulate cortex, hippocampus, occipital and temporal regions, and cerebellar Crus I–II and lobules III, VI, and VIIb (Fig. 3a, left). In the right hemisphere, elevated degree was observed in extensive frontal, temporal, parietal, occipital, and cerebellar regions, notably the precentral and superior frontal gyri, insula, cingulate cortex, hippocampus, and cerebellar Crus I–II and lobules III–VIII (Fig. 3a, right).

**Figure 3.**
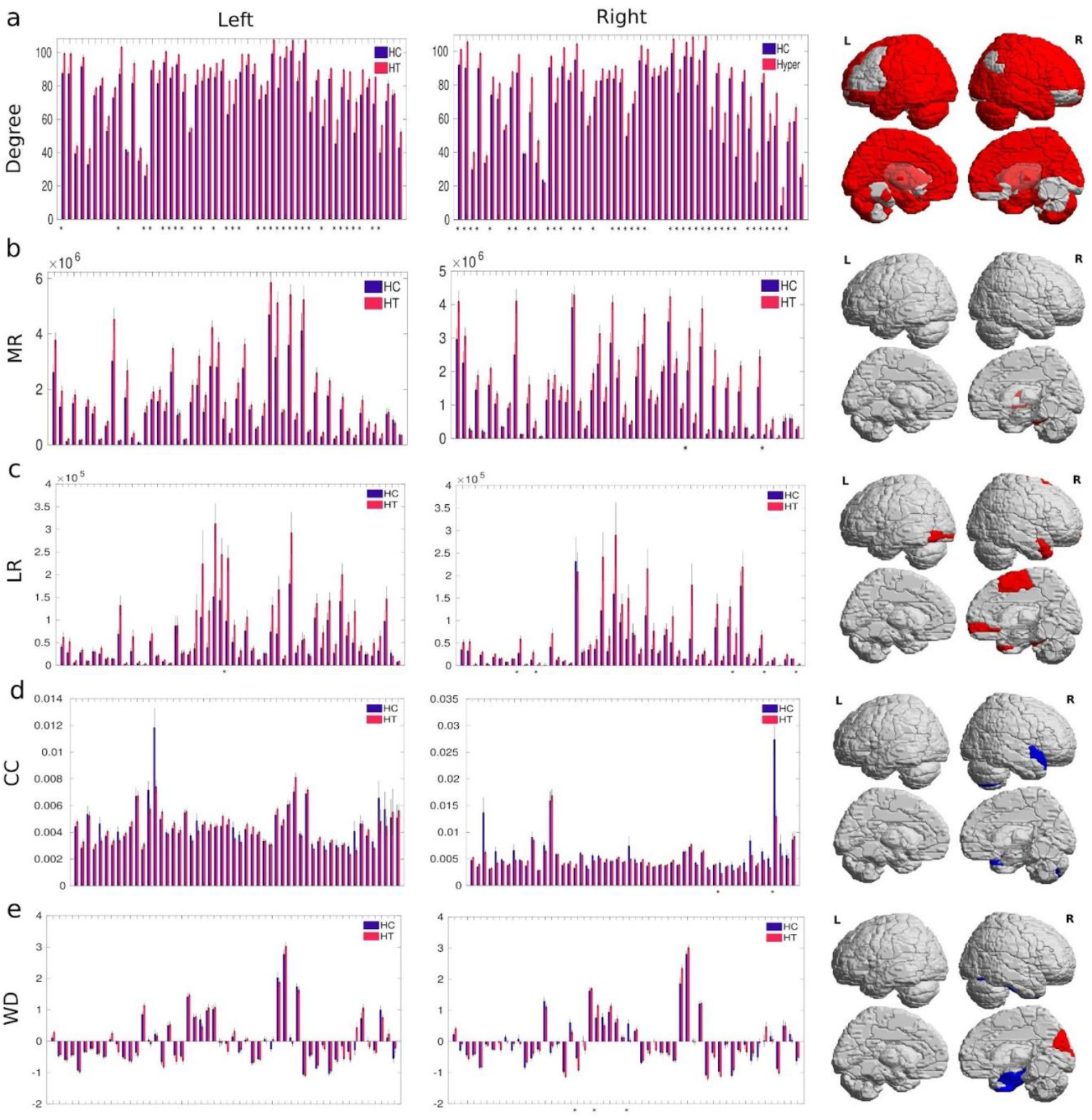
Node level differences in network topology between HC and HT groups: (a) Regional degree, (b) region wise middle range (MR), (c) region wise long range(LR) connection strength, (d) Clustering coefficients (CC), and (e) Within-module degree z-scores (WD), compared between groups across left hemisphere (left) and right hemisphere (middle) brain regions. Bars represent group means for each region, with error bars indicating standard error of the mean (SEM). Brain ROIs (right panel) are color-coded to reflect significant differences: red indicates higher values and blue indicates lower values in the HT group relative to HC. Asterisks (*) denote regions with statistically significant differences between groups after FDR correction (FDR-corrected p < 0.05).

For connection strength, mid-range (MR) connectivity increased in the right putamen and cerebellar lobule IV–V (Fig. 3b), while long-range (LR) connectivity was elevated in the left inferior occipital gyrus and in the right supplementary motor area, middle orbital frontal gyrus, middle temporal pole, and cerebellar lobules IV–V and IX (Fig. 3c). No FDR-corrected differences were found for short-range (SR) connections.

For CC, significantly reduced local segregation was observed in the right superior temporal pole, and cerebellum lobule VII b (Fig. 3d).

WD was significantly lower in HT the right parahippocampal gyrus and fusiform gyrus, and significantly higher in the right cuneus (Fig. 3e). No FDR corrected differences between groups were found for PC and BW.

### Reorganization of hub architecture in HT

We consensus averaging the SC over each group to investigate hub-level reorganization. Classification of nodes based on our hub detection algorithm (see Method, Fig S1a) revealed a shift in hub roles, with regions such as the left putamen and left caudate transitioning from provincial to connector hubs in the HT group (Fig. 4). Subject level hub architecture analysis added in Fig S1b. Both groups showed hubs primarily within cortical and subcortical association areas; however, the HT exhibited a higher prevalence of connector hubs (red nodes), particularly in frontoparietal and limbic regions, suggesting altered inter-modular integration. Node size represents the number of subjects in which a given region was identified as a hub, indicating greater consistency of hub organization in the HT group.

**Figure 4.**
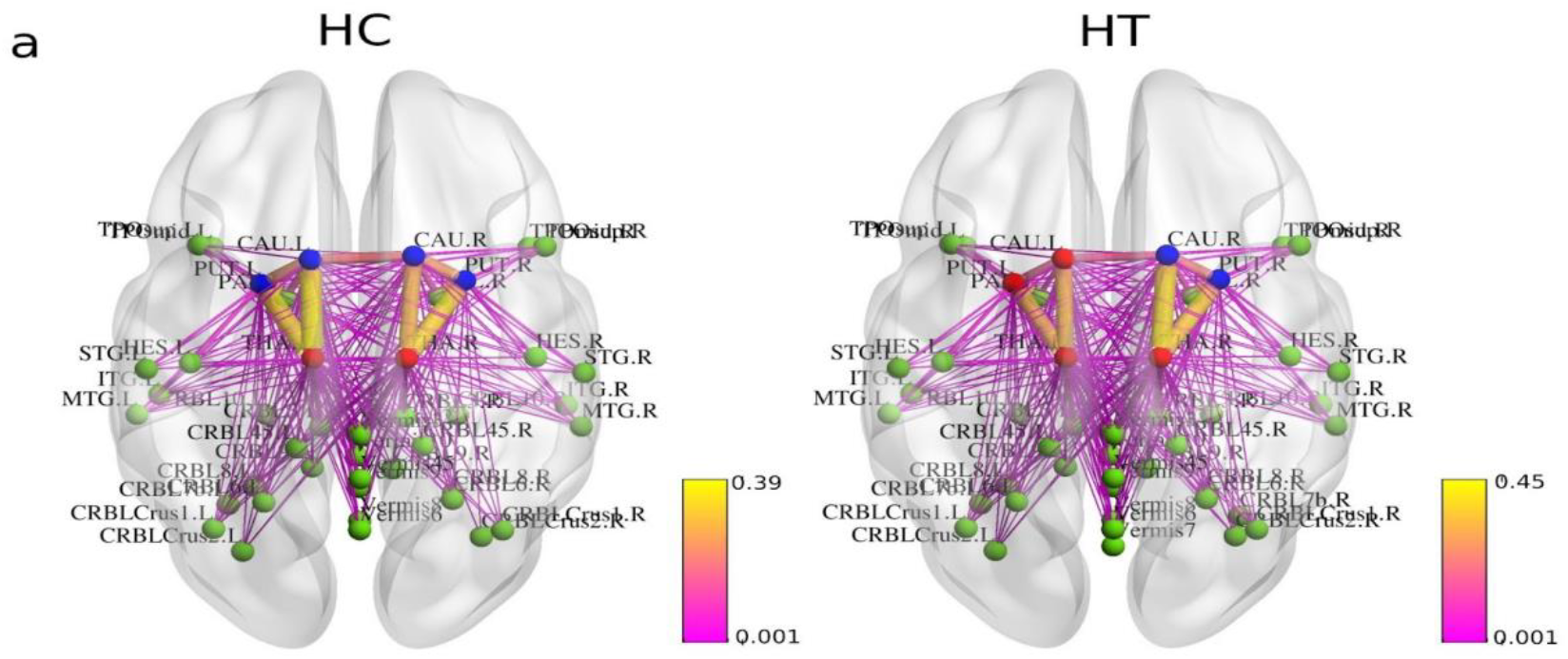
Alterations in hub architecture and node roles in average HC and HT groups: 3D brain renderings showing the spatial distribution of hub nodes. Connector hubs (red), provincial hubs (blue), and non-hubs (green) are visualized for Control and Hyper groups, with major structural connections (pink lines). Provincial hub of Control group becomes connector hubs in Hyper group e.g left putamen and left caudate.

**Figure 4.**
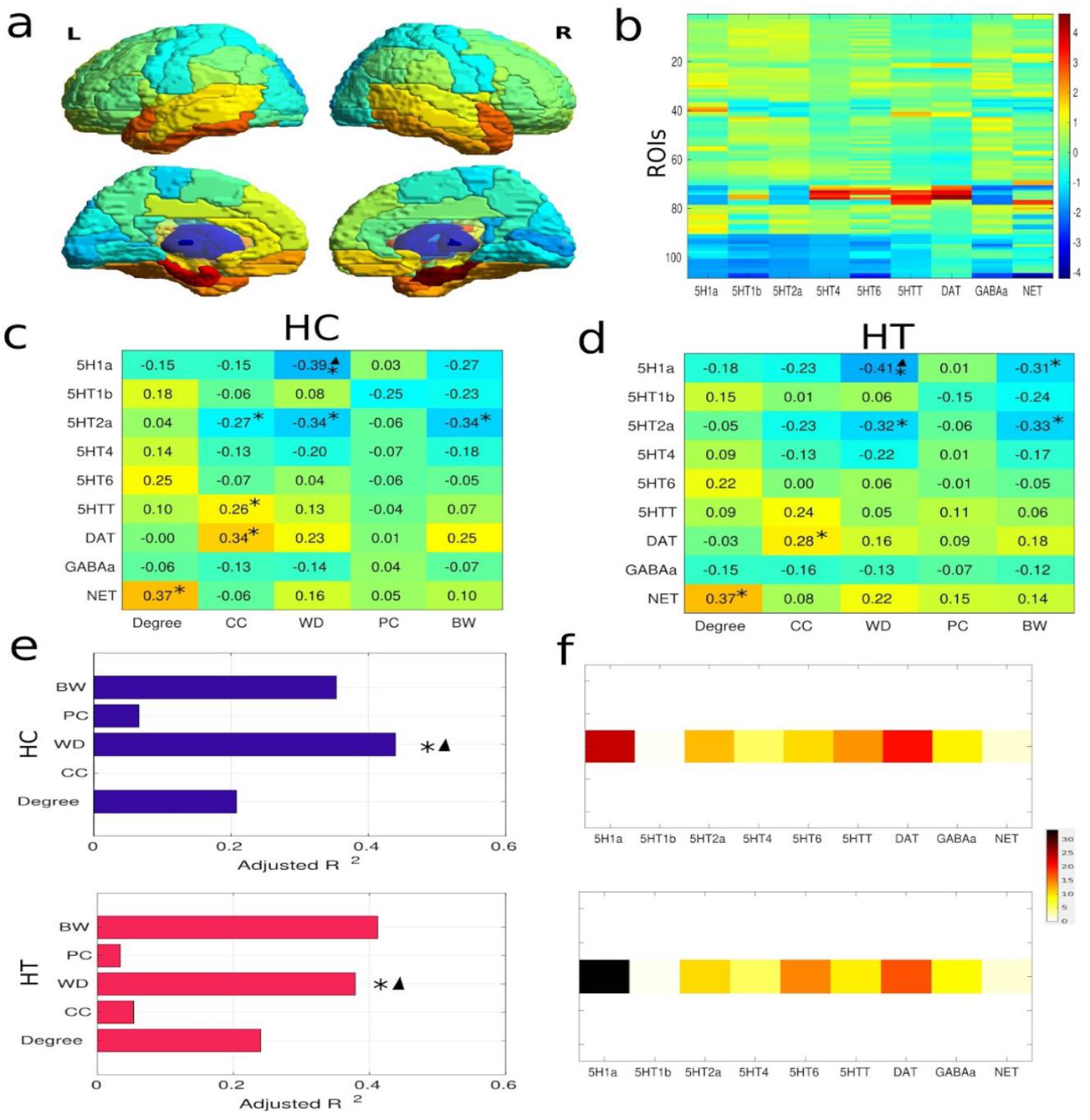
Associations between neurotransmitter receptor densities and network topology in average control and hyper groups: (a) PET maps for 9 different neurotransmitter receptors and transporters were z-scored and collated into a single neurotransmitter receptor atlas. (b) Heatmap showing z-scored normative neurotransmitter receptor and transporter densities across 116 brain regions (ROIs) for 9 neurotransmitters (columns). Warmer colors indicate higher relative receptor density (after z-scoring across regions). (c, d) Group-level Spearman correlations between neurotransmitter receptor density and graph metrics for HC and HT respectively. Asterisks (*) indicate significance based on permutation testing (FDR-corrected p < 0.05). Triangles (▴) indicate significance based on spin permutation testing (p < 0.05). Warmer colors indicate stronger positive associations. (e) We fit multiple linear regression models to predict different network measures from neurotransmitter receptor distributions separately for each group (Control: upper panel; Hyper: lower panel). Adjusted R^2^ values represent the explanatory power of neurotransmitter densities in predicting each network metric. Asterisks indicate significant models (FDR corrected p_perm_< 0.05) while triangles (▴) indicate significance based on spin permutation tests (p_spin_<0.05). (f) Dominance analysis distributes the fit of the model across input variables such that the contribution of each variable can be assessed and compared to other input variables (HC: upper panel; HT: lower panel). The percent contribution of each input variable is defined as the variable’s dominance normalized by the total fit (Adjusted R^2^) of the model. Note that dominance analysis is not applied to the input variables of non-significant models.

### 3.3 Association between network metrics and cognitive and clinical variables

BMI of HT was positively associated with modularity (R^2^ = 0.21, coefficient = 1.12, p = 0.01), suggesting that lower BMI may be linked to decreased modularity of the structural network in HT (Tab. 2).

Free thyroxine (FT4) of HT was positively associated with characteristic path length (R^2^ = 0.16, coefficient = 0.392, p = 0.03) and long range (LR) connection strength (R^2^ = 0.14, coefficient = 0.376, p = 0.04), suggesting that higher FT4 may be linked to increased characteristic path length and LR connections strength of the structural network. We do not find any significant association with other cognitive scores and global network measures. However, local measure roi-wise MR connection strength and degree of right cerebellar lobule IV-V was associated with increased error rate in perceptuomotor function tasks (Nahor benson test) (R^2^=0.143, coefficient=3.706, p<0.05 for MR ; R^2^=0.119 coefficient=1.82, p<0.05) (Tab. 3).

### 3.4 Interdependence of topological measures on neurotransmitter density

Normative PET-derived neurotransmitter density maps from Hansen et al. (2022) were parcellated into 116 regions using the AAL atlas (Tzourio-Mazoyer et al., 2002; Fig. 4a). Z-scored receptor and transporter density maps for nine neurotransmitters showed marked regional heterogeneity (Fig. 4b). Spearman correlations between receptor densities and graph metrics were computed separately for healthy controls (HC) and hyperthyroid (HT) groups (Figs. 4c–d).

Multiple linear regression models predicting network-level metrics from receptor densities were evaluated against spin-permuted null models (10,000 repetitions). Only the model for within-module degree (WD) was significant after FDR correction (P_spin_ < 0.05, one-sided), showing moderate fits (Adjusted R^2^ = 0.45 for HC; 0.38 for HT; Fig. 4e). Dominance analysis revealed that 5-HT_1_A (25% HC; 33% HT) and DAT (20% HC; 17% HT) contributed most to explaining WD variance (Fig. 4f).

Serum TSH levels were significantly lower in HT compared to HC (Fig. 5a). Neurotransmitter-weighted WD distributions for 5-HT_1_A and 5-HTT (Fig. 5b) showed a significant association with TSH (p = 0.04), explaining moderate variance (R^2^ = 0.52; Adjusted R^2^ = 0.30). In the HT group, predicted versus observed TSH values (Fig. 5c) indicated that 5-HTT was a significant positive predictor (Coeff. = 9.28, p = 0.03), while 5-HT_2_A, DAT, GABA, and NET showed moderate positive trends without reaching significance (Fig. 5d).

**Figure 5.**
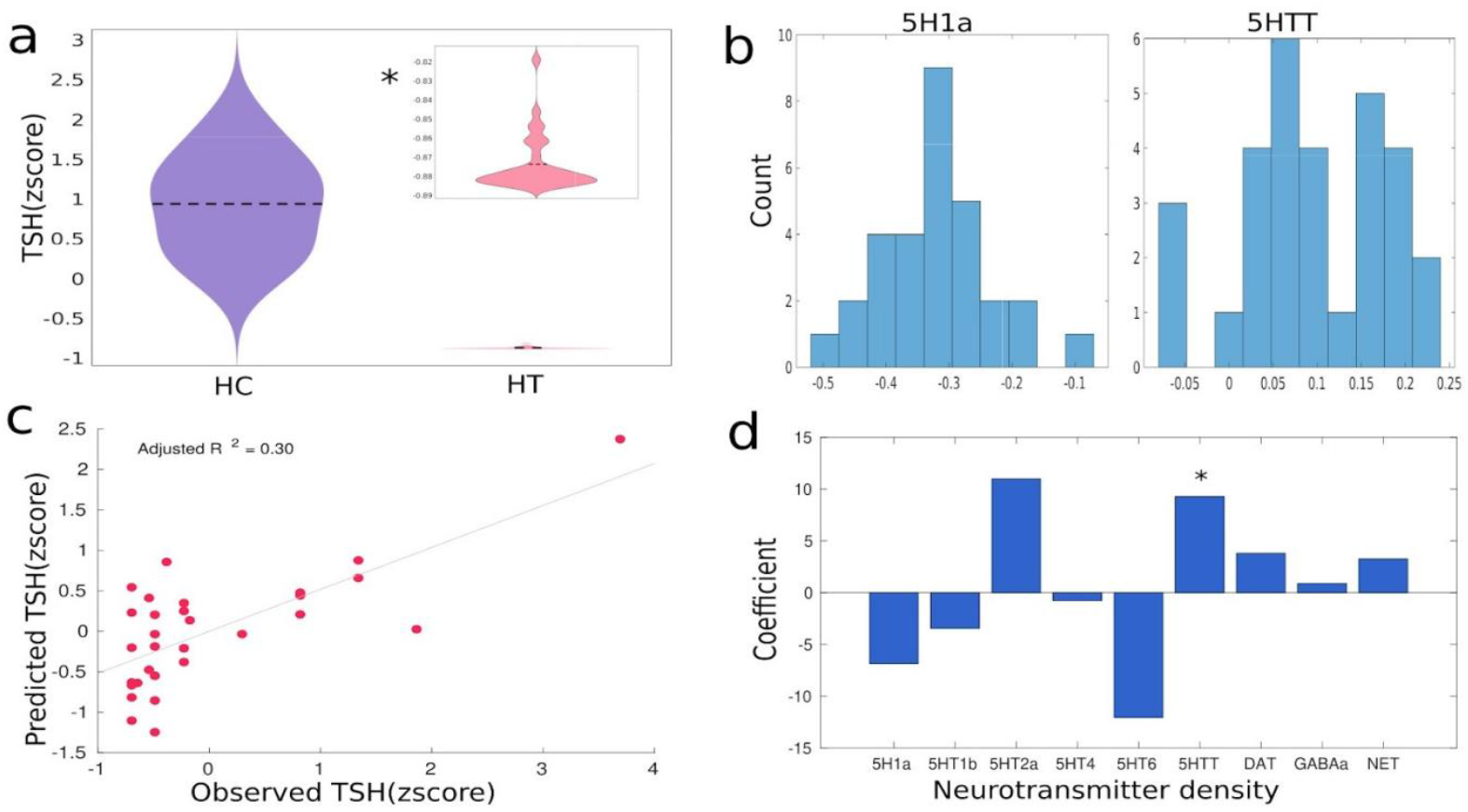
Association between serum TSH levels, neurotransmitter weighted structural network topology. (a) Violin plot of TSH (z-score) for the HC and HT groups. Asterisks (*) indicate significant differences between groups. The inset violin plot illustrates the distribution of TSH values within the HT group. (b) Distribution of neurotransmitter weighted WD over subjects (left panel 5H1a and right panel 5HTT-weighted WD). (c) Multivariate regression model predicting TSH levels in the HT group using neurotransmitter-weighted within-module degree z-scores (WD) as predictors. (d) Standardized regression coefficients for each neurotransmitter-weighted WD predictor; among them, the 5-HTT-weighted WD z-score shows a significant association with TSH (p <0.05).

## 4. Discussion

This study characterized alterations in the structural brain network topology associated with hyperthyroidism and examined their links to cognitive function, clinical parameters, and molecular architecture. Individuals with hyperthyroidism showed maladaptive structural reorganization, marked by increased mid- and long-range interhemispheric coupling and disrupted network topology, most prominently in the right hemisphere. These topological changes were functionally relevant, as global metrics correlated with clinical and cognitive variables such as fT4, BMI, and perceptuo-motor scores. Spatial correspondence analyses revealed that dopaminergic, serotonergic, and noradrenergic markers—particularly DAT, 5-HT2A, and NET-were associated with network alterations. Moreover, subject-level neurotransmitter-weighted features significantly predicted TSH levels, linking hormonal imbalance to molecular–connectomic reorganization. This integrative framework highlights molecular contributions to macroscale network changes in hyperthyroidism.

Our structural network analysis revealed a significant reduction in modularity indicative of decreased network segregation. A highly modular architecture is a cornerstone of efficient brain function, allowing for specialized, parallel processing of information within distinct functional communities at a low metabolic and wiring cost (Van Den Heuvel & Sporns, 2013; Sporns & Betzel, 2016). In the hyperthyroid group, this reduction in modularity coincided with a significant increase in characteristic path length, reflecting longer routes for information transfer between brain regions. Together, these findings point to a compromise in the brain’s ability to sustain functionally specialized modules, a phenomenon linked to reduced processing efficiency and cognitive impairment in other neurological and psychiatric conditions (Fornito et al., 2015).

Closer inspection revealed that these global alterations were driven by strengthened medium- and long-range hemispheric connections. Paradoxically, while such long-range connections might be expected to reduce path length and promote integration (Betzel & Bassett, 2018), our data showed no improvement in global efficiency. Instead, the proliferation of extended connections coincided with reduced segregation and longer communication routes, implying redundant or misrouted connectivity. This inefficiency may also reflect the elevated metabolic demands of hyperthyroidism (Lönn et al., 1998), as longer connections are energetically more costly.

This conception is further supported by the clinical correlation with global network measures. In the hyperthyroid group, modularity significantly and positively predicted BMI. The concurrent reduction in modularity and BMI in HT suggests that the observed network reorganization may be both topologically inefficient and metabolically costly (Ríos-Prego et al., 2019). Specifically, the increase in redundant, spatially extended connections may impose greater energetic demands on the brain’s architecture, reflecting and possibly contributing to the systemic increase in the metabolic rate that characterizes hyperthyroid states. In parallel, free thyroxine levels were positively associated with both characteristic path length and long-range connection strength, indicating that excess fT4 drives the formation of extended connections but in a way that paradoxically reduces communication efficiency. FT4, the biologically active thyroid hormone, is known to modulate neuronal excitability, synaptic plasticity, and myelination (Bernal, 2007; Bauer et al., 2008). Chronic exposure to elevated fT4 may enhance conduction velocity and white matter organization, yet in this context it appears to promote maladaptive network wiring.

Hence, in hyperthyroidism, we observed a maladaptive reorganization of structural brain networks, marked by increased medium- and long-range hemispheric connection strength, reduced modularity, and longer characteristic path length. These inefficient wiring patterns may be driven by excessive fT4 and are further linked to reduced BMI (He et al., 2021). The enhanced long-range connections strength likely impose higher energetic demands for information transfer, suggesting that increased resting metabolic rate is mirrored in the brain’s network architecture.

Regionally, most brain areas exhibited increased white matter connection strength in hyperthyroidism, with more pronounced effects in the right hemisphere. The pattern of increased nodal degree indicates focal proliferation of white matter connections, aligning with reports of elevated cortical thickness in right-hemisphere regions of hyperthyroid patients correlated with thyroid hormone levels (Kumar et al., 2024). Notably, the right putamen and cerebellar lobule IV–V showed increased degree and mid-range connection strength, implicating basal ganglia and cerebellar circuits involved in motor control (Houk & Wise, 1995; Bradshaw & Sheppard, 2000), cognitive regulation (Botvinick & Braver, 2015), and coordination (Peterburs & Desmond, 2016). The association of cerebellar lobule IV–V connectivity with psychomotor dysfunction supports a link to agitation, anxiety, and arousal in hyperthyroidism (Fahrenfort et al., 2000). Reduced local segregation in the right superior temporal pole and cerebellum (lobule VIIb) further underscores diminished specialization.

The right parahippocampal gyrus, a region central to contextual and spatial memory (Baker et al., 2018), exhibited a significant increase in degree alongside a significant reduction in WD in our hyperthyroid cohort, pointing to regional disconnection or diminished intra-modular hubness despite excess connection strength within memory circuits. Additionally, analysis revealing alterations in several brain regions critically involved in memory (such as right middle temporal pole, right fusiform gyrus, right thalamus) showed a significantly higher degree in hyperthyroidism. Although, we didn’t observe any significant correlation, however, these findings may provide a neuroanatomical substrate for the memory impairments in hyperthyroidism (Vogel et al., 2007; Yudiarto et al., 2006, Sahin et al., 2023).

Together, these results highlight hemispheric asymmetry in hyperthyroidism, aligning with a broader literature on lateralized network pathology in neuropsychiatric disorders such as autism (Floris et al., 2021), schizophrenia (Lohr & Caligiuri, 1997), and other psychiatric disorders (Huang et al., 2021). Hub analysis revealed reconfiguration, with the left putamen emerging as a connector hub-potentially reflecting compensatory adaptation to right-lateralized connectivity changes.

Integrating PET-derived neurotransmitter receptor maps, we found that nodal-level features, particularly WD and betweenness centrality (BW), were shaped by underlying molecular architecture. Serotonin (5-HT1A, 5-HT2A) and dopamine (DAT) receptor distributions were key predictors of network topology. Regions with higher 5-HT1A density showed lower WD, indicating reduced local hubness, and dominance analysis confirmed that 5-HT1A and DAT contributed most to WD variance. The right parahippocampal gyrus-critical for contextual and spatial memory (Aminoff et al., 2013; Bohbot et al., 2015)—had significantly reduced WD in the hyperthyroid group. Given evidence that hippocampal 5-HT1A activation supports spatial learning (Glikmann-Johnston et al., 2015; Gerdey & Masseck, 2023), serotonergic modulation may underlie diminished hubness and memory deficits in hyperthyroidism. However, a direct relationship between topological reorganization and memory related problems were not observed which might be related to the specificity of the cognitive test included in our study.

Additionally, it has been suggested that 5HT1a and 5HT2a play an important role in plasma level TSH (Sullo et al., 2011) secretion. In particular, an inhibitory control of TSH plasma level by 5HT has been suggested in earlier animal studies (Mattila & Männistö, 1981 ; Di Renzo et al., 1979). Similarly, we also observed a positive association between TSH and 5HTT weighted WD, suggesting a possible role in its indirect regulation at plasma level in hyperthyroidism. Thus, association between topological network alterations and normative spatial neurotransmitter/transporter densities provides crucial insights into the neurobiological mechanisms underlying hyperthyroidism.

## 5. Limitations and future work

Although, we observed significant differences in topological SC but small sample size limits the generalization of the results. Additionally, our study is a cross-sectional design that restrains us from making speculations regarding the involvement of topological measures in disease progression or modification.

We included the neurotransmitter density maps from the open access healthy individual brains data, which is not having any specificity to hyperthyroidism. Individual neurotransmitter density maps could have provided the hyperthyroid specific profiling. Although the topological SC measures correlated with the cognitive impairments in hyperthyroidism, adding functional connectivity based topological measures in future study could overcome a direct relationship with cognitive measures. Finally, the moderate predictability of neurotransmitter weighted network measures for TSH hormone could be due to small sample size or due to non cohort specific neurotransmitter measures. A future longitudinal study including multimodal approach in a larger cohort may warrant the validation of our findings.

## 6. Conclusion

This study demonstrates that overt hyperthyroidism is linked to systematic reorganization of the structural connectome, with pronounced right-hemispheric effects suggesting hormone-driven maladaptive neural plasticity. Associations between clinical parameters (fT4, BMI) and network measures indicate that energetically costly long-range connections may reduce segregation and efficiency, consistent with elevated metabolic demand. Enhanced mid-range connectivity and increased degree in cerebellar lobule IV–V were related to psychomotor dysfunctions, implicating excessive cerebellar connectivity in agitation and anxiety. The spatial distribution of local hubness is also aligned with neurotransmitter systems (5-HT1A, DAT, NET) involved in regulating the hypothalamic–pituitary–thyroid axis. Overall, hyperthyroidism disrupts structural network organization, highlighting the need for larger longitudinal studies to confirm these findings.

## Abbreviations

HT: Hyperthyroidism
HC: Healthy control
BMI: Body mass index
TSH: Thyroid-stimulating hormone
T3: Triiodothyronine
FT4: Free thyroxine
GABA: Gamma aminobutyric acid
DAT: Dopamin Transporter
DWI: Diffusion weighted imaging
PGIBBD: Postgraduate Institute Battery of Brain Dysfunction
SR: Short range connection
MR: Middle range connection
LR: Long range connection
MMSE: Mini Mental State Examination
P/K: Ratio of Pass-a-long and Koh’s test

## Author Contributions

PC contributed to study conception, data analysis, statistical analysis, writing and editing the manuscript. NU contributed to study conception, data analysis, writing and editing the manuscript. PK contributed to data collection, writing and editing the manuscript. TS contributed in recruitment and clinical assessment of thyroid patients and study design, review and editing the manuscript. SS contributed to data analysis, statistical analysis, and editing the manuscript.. SK contributed to problem definition, study design, and review and editing of the manuscript. PR was contributing study concepts and study design, reviewing and editing the manuscript. MD in MRI radiologist, prepared the patient’s report, and reviewed the manuscript. MK contributed to data collection, data analysis, writing and editing the manuscript.

## Acknowledgments

This work was supported by DRDO R&D Project No. INM 311 (4.1). Additionally, I am grateful to DRDO for providing financial support as a research fellow grant while conducting the research activity.

