## Supplementary material for "Right-Lateralized Maladaptive Topological Reorganization in Hyperthyroidism"

### Authors contributed equivalently

Dr. Priyanka Chakraborty

Dr. Neeraj Upadhyay

Dr. Mukesh Kumar

#### Material and method

##### Subjects

Drug-naive thirty hyperthyroid patients (age, 34.63± 8.99 years; 10 males; 20 female) and twenty-eight healthy controls (HC) (age 32.82± 9.86 years; 9 males; 19 female) were recruited for this study. Hyperthyroid patients first time diagnosed with clinical symptoms such as palpitation, trembling, loss of weight and heat intolerance from days to weeks. Patients were recruited from the Thyroid Research Centre, Institute of Nuclear Medicine and Allied Sciences (INMAS) and HC were recruited from the local community. Participants with a history of trauma, psychiatric illness, substance abuse, pregnancy, or contraindication with MRI were excluded.

##### Clinical assessment/Thyroid Hormone tests:

Blood samples were collected from both hyperthyroid and healthy subjects in the morning between 8.30-10 am with empty stomach under fasting conditions to minimize the influence of diurnal variations. The serum thyroid indices, [fT4 (free thyroxine), fT3 (free triiodothyronine) and TSH (thyroid-stimulating hormone)], were measured using an Electro Chemiluminescence Immuno Assay Kit (Elecsys 2010; Roche Diagnostics, Mannheim, Germany).

Thyroid hormones of the control subjects were within normal ranges (TSH = 0.27-4.2 µIU/ml, fT4 = 12.0-22.0 pmol/l, and fT3=2.6-6.8 pmol/l). While, hyperthyroid patients, the suppressed TSH (.01±.019µIU/ml), elevated fT4 (42.91±18.58 pmol/liter) and fT3 (15.46±5.56 pmol/liter) levels were observed. (Details in Table 1).

**Neuropsychological Data**

All the patients underwent through the Mini Mental State Examination (MMSE) (Folstein et al., 1975) and a battery of neuropsychological tests 'Postgraduate Institute Battery of Brain Dysfunction' (PGIBBD) (Pershad D, 1990), before the MRI scan. The PGIBBD battery, which is used to assess varieties of cognitive functions including five major tests: PGI Memory Scale (PGIMS), Revised Bhatia Short Battery of Performance Tests of Intelligence (BSR-R), Verbal Adult Intelligence Scale (VAIS), Bender Gestalt test (BGT) and Nahar-Benson test (NBT)(Pershad D, 1990). PGIMS (consisting of ten subtests) was selected to assess the long term episodic memory, recent episodic memory, mental balance, working memory span (forward and backward), delayed and immediate recall memory, immediate recall of semantically related word pairs, immediate recall of arbitrarily related word pairs, visual retention and recognition of the objects memory. In PGIMS subtests, higher scores indicated better performance.

Executive function performance quotient was measured by BSR-R subtests, performance quotient score includes (Koh's Block Design (KBD) and Pass-a-Long test (PALT)). Higher scores in KBD and PALT showed better performance. Motor function was evaluated using the Nahar–Benson Test (NBT), while visuospatial function was assessed with the Bender Gestalt Test (BGT). Higher dysfunction scores (Dys) on these tests indicated poorer performance as well. The scoring of all the neuropsychological tests was done as per the procedure described in detail elsewhere (Pershad and Verma, 1990).

**Structural data acquisition:**

The study was carried out on a 3T whole-body MR system (Magnetom Skyra; Siemens, Erlangen, Germany) with 20 channel head and neck coil and a 45mT/m actively shielded gradient system. Diffusion-weighted data was acquired using a single-shot echo-planar dual SE sequence, on parameters: 5 b-values = 0, 500, 1000, 1500 and 2000 s/mm2 with 30 diffusion direction, slice thickness = 4.5 mm with no inter-slice gap, number of slices = 30, FOV = 230 mm × 230 mm, matrix size = 128 × 128, spatial resolution = 1.797 mm X 1.797 mm X 4.5 mm, flip angle 90°, TR/TE = 7300/138 ms, number of excitations (NEX) =1.

To rule out any structural abnormalities, routine T2-weighted and T1-weighted images were acquired using the following parameters: for T2-weighted imaging, TR/TE = 5600/100 ms, slice thickness = 4.0 mm, 25 slices, number of excitations (NEX) = 2, matrix size = 312 × 512, and field of view (FOV) = 220 mm; for T1-weighted imaging, TR/TE = 1900/2.49 ms, slice thickness = 0.9 mm, 160 slices, matrix size = 256 × 256, and FOV = 240 × 240 $mm^2$.

**DTI preprocessing**

The diffusion MRI data were processed locally using our pre-processing pipeline, which involved using MRtrix3 ((Tournier et al.2019), https://github.com/MRtrix3/mrtrix3), FSL (http://www.fmrib.ox.ac.uk/fsl), and ANTs (http://stnava.github.io/ANTs/). The main preprocessing steps included denoising, (Veraart et al. 2016), removal of Gibbs ringing artifacts (Kellner et al. 2016). Susceptibility distortion correction of DKI was performed using the Synb0-DISCO algorithm (Schilling et al., 2019, 2020). Bias correction using the MRtrix with ANTs (Tustison et al. 2010). The T1-weighted mask was registered to the diffusion space to generate a brain mask in diffusion space. We applied multi-shell multi-tissue constrained spherical deconvolution (MSMT-CSD) to determine the orientation of the fibers [fibre orientation distribution {FOD}] in each voxel. After that, we normalised the global intensity such that the FODs were similar among subjects. To create a tissue-segmented picture, we employed anatomically constrained tractography (ACT) (Smith et al. 2012). In order to simplify seeding, we made a mask of the grey matter/white matter border. Lastly, we produced 10 million tracts using probabilistic tractography. In order to identify a subset of streamlines with streamline densities that were closer to fibre densities, the tractogram was further filtered (Smith et al. 2015). Each step was visually assessed and edited by research personnel.

**Structural connectivity and fiber tract length**

A subject-specific whole-brain connectome was created, calculating the fiber density between each pair among the 116 ROIs, using the Automated anatomical labelling (AAL) atlas (Tzourio-Mazoyer et al., 2002). We were then able to quantify the number of streamlines that started from one ROI and ended on one of the other ROIs as well. The physical length of the fiber in millimeters between each pair of ROIs was calculated, with a maximum length of 250 mm. For each subject, a tract length (TL) matrix was created based on the AAL parcellation. Then we defined short (SR), middle (MR), and long (LR) connections based on TL, setting two thresholds, one at 70 mm and another at 140 mm (Chakraborty et al., 2025). A group consensus binary network that maintains the density and edge-length distributions of the individual connectomes was built for both over group analysis and individual subject wise analysis (Betzel et al., 2019). This process more accurately reflects significant organisational characteristics of subject-level networks (i.e., thresholding based on whether an edge is noticed in a fraction of subjects). By averaging the log-transformed streamline count of non-zero edges among participants, weights were assigned to the group consensus network's edges. After that, edge weights were scaled to values ranging from 0 to 1.

**Graph theoretical analysis**

(i) *Degree:* The degree of a node in a network is the number of connections (edge) attached to it (here white matter connection). It represents how many direct neighbors a node has in an unweighted network. The mean of all node degrees in a network gives the average degree of the network.

(ii) *Modularity:* Modularity assesses the degree to which a network may be separated into different communities with dense intra-module connections and sparse inter-module connections. High modularity denotes robust community structure and functional

specialization (Newman, 2006). Modularity gives network resilience and adaptability, measuring the degree of segregation. Communities are subgroups of densely interconnected nodes sparsely connected with the rest of the network.
*(iii) Clustering coefficient:* Local clustering coefficient represents the abundance of connected triangles in a network (Watts & Strogatz, 1998).

(iv) *Transitivity:* This referred to as the global clustering coefficient, measures the likelihood that a node's neighbours are connected as well, indicating how common closely connected clusters are in the network. According to (Humphries & Gurney, 2008), it is calculated as the graph's triangle to triplet ratio. Stronger modular segregation and local connectivity are indicated by higher transitivity.
(v) Characteristic *path length:* The characteristic path length is the average shortest path length between all pairs of nodes in the network, representing the average distance between nodes (Watts & Strogatz, 1998). It is an indicator of global integration, inversely related to global efficiency. Lower path length implies faster communication across the network. We use tract length matrix for calculation of char. path length.

(vi) *Global efficiency* The average inverse shortest path length between every pair of nodes in the graph is known as global efficiency, and it serves as a gauge of network integration. According to Latora and Marchiori (2001), it shows how well information is shared across the network. (Latora & Marchiori, 2001)

(vii) Within-module degree z-score (WD): The within-module degree z-score quantifies how well-connected a node is to other nodes within its own community. A positive z-score indicates that the node is more connected within its module compared to the average node in that module, whereas a negative z-score suggests fewer connections.

(viii) Participation coefficient (PC) A graph's modular structure allows for the computation of each node's participation coefficient . (Guimerà & Nunes Amaral, 2005). PC measures how well distributed the links of a node are to other modules. A high participation coefficient indicates that a node is a connector hub involved in multiple modules, whereas a low value suggests that the node's connections are confined within its own module. .

(ix) Betweenness centrality Kintali (2008): Betweenness centrality quantifies the number of shortest paths that pass through a given node, indicating the node's control over information flow in the network. High betweenness centrality nodes act as bridges and are critical for global communication.

**Neurotransmitter maps from positron emission tomography**

Receptor densities for 19 receptors and transporters across nine neurotransmitter systems were estimated using PET tracer studies, as recently made available by Hansen and colleagues (Hansen, Shafiei, Markello, et al., 2022). We estimated nine receptor and transporter densities using PET tracer studies. These systems include dopamine (DAT), norepinephrine (NET), serotonin (5-HT1a, 5-HT1b, 5-HT2a, 5-HT4, 5-HT6, 5-HTT), GABA (GABAa) on the basis of earlier studies on hyperthyroidism (Napoli et al., 2001; Rastogi & Singhal, 1976; Wiens &

Trudeau, 2006b). We extracted regional neurotransmitter densities for these 9 receptors and transporters using the AAL atlas, then z-scored each map before compiling them into a region × receptor matrix of relative densities.

**Statistical analysis**

Statistical analysis was performed on IBM SPSS (version 27.0, SPSS Inc, Chicago, IL, USA) statistics. Demographic and clinical parameters were assessed by independent sample t-tests (two-tailed), and gender was compared using the Pearson Chi-square test. Analysis of covariance (ANCOVA) was used to compare neuropsychological scores between the two groups, with age and gender as covariates. All the differences in demographic, clinical and neuropsychological measures were reported at $p<0.05$.

To assess group differences in brain network organization, we compared each graph-theoretical metric between the HT and HC groups. First, we assessed the normality of the data implementing Shapiro-Wilkinson's test. Then a t test was applied to evaluate global as well as regional differences for each metric independently. To account for multiple comparisons across regions and metrics, resulting p-values were subjected to false discovery rate (FDR) correction (Benjamini & Hochberg, 1995). Furthermore, to ensure statistical robustness and control for potential biases, we used permutation testing (n = 10,000 iterations). This involved randomly shuffling group labels and re-evaluating test statistics to generate a null distribution. For analyses involving receptor maps, we implemented non-parametric spin permutation tests (Alexander-Bloch et al., 2018; Markello & Mišić, 2021) to account for the spatial autocorrelation inherent in brain data.

We employed multiple linear regression models for various predictive analyses. Age, gender, and years of education were treated as covariates in all analyses and hence regress out them before fitting the model. Model performance was evaluated using adjusted $R^2$. Generalizability of the model was evaluated using 10,000 permutations and 10,000 spin permutations for neurotransmitter maps.

To assess the relative importance of individual predictors in explaining variance in the outcome variable, we employed dominance analysis (Azen, R. & Budescu, D. V). This technique decomposes the total model $R^2$ by systematically evaluating all possible subset models, allowing for a rank-ordering of predictors based on their average additional contribution across models. For each predictor, the general dominance weight was computed as the average increase in $R^2$ when the predictor was added to all subset models that excluded it.

**Result**

Supplementary Table 1. Demographic, clinical and neuropsychological characteristics of hyperthyroid, as compared to healthy controls.

| **Characteristics** | **Controls (N = 28)** | **Hyperthyroid (N = 30)** | **p-value** |
|---|---|---|---|
| **Memory scale** | | | |
| Long term episodic memory | 4.82 ± 0.61 | 4.67±0.66 | .77 |
| Recent episodic memory | 4.96± 0.19 | 4.83±0.46 | .92 |
| Delayed Recall Memory | 4.86±0.36 | 4.57±0.94 | .28 |
| Immediate Recall Memory | 4.79±0.42 | 4.63±0.81 | .88 |
| Immediate recall of semantically related word pairs Memory | 4.93±0.26 | 4.30±1.12 | .05 |
| Immediate recall of arbitrarily related word pairs Memory | 4.54±0.84 | 4.03±1.30 | .41 |

| | | | |
|---|---|---|---|
| Visual Retention Memory | 4.79±0.42 | 4.23±1.19 | .19 |
| **Performance Tests of Intelligence** | | | |
| Performance quotient | 113.88±19.65 | 103.18±19.61 | .21 |
| P/K*100 | 135.39±82.71 | 154.04±100.08 | .89 |

Mean and standard deviation (SD) for demographic, blood hormone levels, neuropsychological tests score.

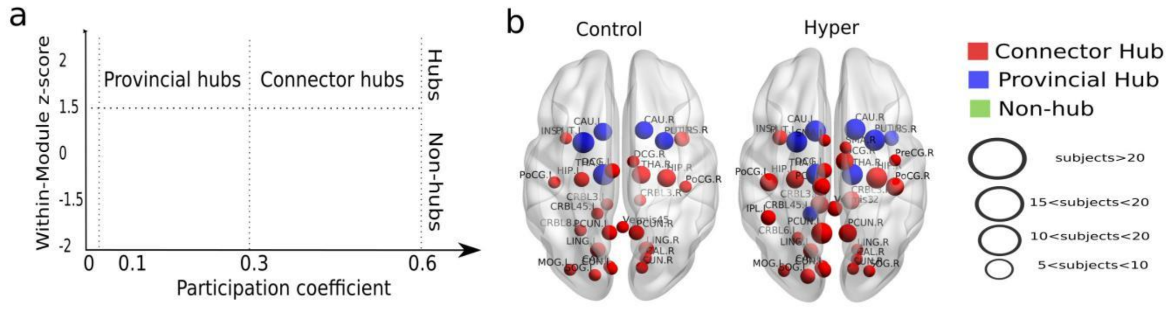


Figure S1 (a) Nodes are categorized based on their subject-level values of within-module degree z-score and participation coefficient, plotted using the standard cartographic framework. (b) Brain surface renderings show the spatial distribution and prevalence of hubs across the Control (left) and Hyperthyroid (right) groups. Node color indicates hub type, and node size represents the number of subjects (as per scale) in which each region was identified as a hub.
